# A secreted LEA-like protein functions as a basement membrane stress protectant

**DOI:** 10.64898/2026.10.23.761354

**Authors:** Bailey A. Brown, Anna L. Lewis, Madison K. Lester, Zhirong Wang, Jonathan D. Hibshman

## Abstract

Organisms have evolved to withstand a wide variety of environmental stresses. Survival requires individual cells to mount appropriate responses to combat molecular damage. In animals, multicellularity introduces an additional challenge: how is tissue integrity preserved? Our knowledge of extracellular tissue-level mechanisms of protection remains limited. Here, we report that a late embryogenesis abundant (LEA)-like protein, DUR-1, is secreted and then accumulates in basement membranes, where it acts to promote tissue integrity and stress survival. DUR-1 is strongly upregulated in the stress-tolerant dauer diapause stage of *C. elegans* and further upregulated upon desiccation. Loss of the protein sensitizes animals to desiccation and osmotic stress. Other LEA proteins have been regarded as strong desicco-protectants, but none have been recognized as components of basement membranes. We show that DUR-1 is important for maintaining the integrity of basement membrane-ensheathed tissues under mechanical stresses associated with drying or osmotic challenge. This work defines a mechanism for extracellular protection and emphasizes the importance of preserving tissue integrity during stress.

## Introduction

Stress is ubiquitous, and organisms must cope with variable and, at times, harsh environmental conditions to survive. Cellular protection and repair mechanisms support survival under a range of cellular stresses. Examples of cellular stress responses include DNA damage response pathways, antioxidants that mitigate oxidative damage, and mechanisms to maintain proteostasis such as the unfolded protein response, autophagy, upregulation of chaperones, and proteasomal quality control [1]. Notably, these responses occur cell-autonomously and, in many cases, are conserved from single-celled organisms to metazoans [2]. Cell-intrinsic stress responses are essential for survival but likely insufficient to stabilize extracellular components and preserve the integrity of tissues and organs [3,4]. Components of the extracellular matrix, like the basement membranes (BM) that surround tissues, are important for maintaining tissue integrity throughout development. However, adaptations for the protection of these components during stress tolerance are not well understood [5–8].

The nematode *Caenorhabditis elegans*, with its relatively simple body plan, is an ideal animal model to study organismal stress tolerance beyond the bounds of a single cell [9]. In response to environmental stressors, *C. elegans* can enter a developmental diapause stage – the dauer larva [10,11]. Dauer larvae are uniquely tolerant of stress, yet the full suite of proteins that contribute to this stress tolerance remains incomplete. One particularly harsh stress is desiccation – the loss of detectable cellular water [12–14]. Desiccation leads to protein aggregation, membrane damage, and DNA damage [13–18]. In *C. elegans*, desiccation survival is exclusive to the dauer stage and requires a period of conditioning at high humidity, during which protectants are upregulated [19–21].

Late embryogenesis abundant (LEA) proteins are one class of desicco-protectants with demonstrated efficacy across multiple organisms, including nematodes, rotifers, brine shrimp, chironomid midges, and many plant species [15,22–27]. Compared to most anhydrobiotic animals that are not well-developed model organisms, *C. elegans* offers a unique opportunity to study the biology of LEA proteins *in vivo*. Previous work demonstrated that *C. elegans* LEA-1 is required for desiccation tolerance and osmotic stress tolerance, contributes to heat shock survival, and reduces protein aggregation [28–30]. *C. elegans* has a second putative LEA protein, DUR-1 (<u>d</u>auer <u>u</u>pregulated), which has been identified as having sequence homology to other LEA proteins [20,22,30]. DUR-1 is a poorly characterized protein. Its defining feature is its upregulation in dauer larvae. One report found that *dur-1(ok1010)* mutants, like *lea-1* mutants, are sensitive to desiccation [20]. Given the important role of LEA-1 in the stress responses of dauer larvae, we aimed to explore the contributions of DUR-1 to dauer stress tolerance. The existence of a pair of LEA proteins in *C. elegans* further provides a strategic opportunity to determine how LEA proteins may interact and potentially cooperate to promote stress tolerance.

Here, we provide a characterization of DUR-1, demonstrating that it is not required for normal lifespan, brood size, or dauer formation. Working with a newly generated null allele and a remade *ok1010* lesion, we report defects in both desiccation survival and osmotic stress survival of *dur-1* mutant dauer larvae. Although LEA-1 and DUR-1 each individually contribute to dauer stress tolerance, we do not see consistent evidence of synergy between the two proteins. This result is consistent with the different localization patterns and structures of these proteins and suggests that LEA-like proteins may be targeted to unique sites to ensure robust tolerance of desiccation and osmotic stress.

Surprisingly, we find that *dur-1* is primarily transcribed in body wall muscle, secreted throughout the body, and accumulates on basement membranes. Loss of *dur-1* led to reduced tissue integrity upon desiccation and exposure to high salt – two stresses that can lead to mechanical strain. Thus, we define DUR-1 as an extracellular protectant that serves to reinforce basement membranes and promote tissue stability. This work defines a new mode of protection during stress – the use of a LEA-like protein as a BM protectant – and it emphasizes the importance of stabilization beyond the plasma membrane of a single cell.

## Results

### DUR-1 is enriched in dauer larvae but does not impact dauer formation

*dur-1* is named based on its <u>d</u>auer <u>u</u>p-regulated expression. We quantified stage-specific expression of the protein with an endogenous fluorescently tagged *dur-1*::mNG reporter (Figure 1a). As expected, fluorescence intensity was highest in dauer larvae. However, fluorescence was still detectable in other larval stages and adults. This suggests that although largely a dauer-specific protein, DUR-1 is also expressed in other stages and may accumulate throughout development.

**Figure 1.**
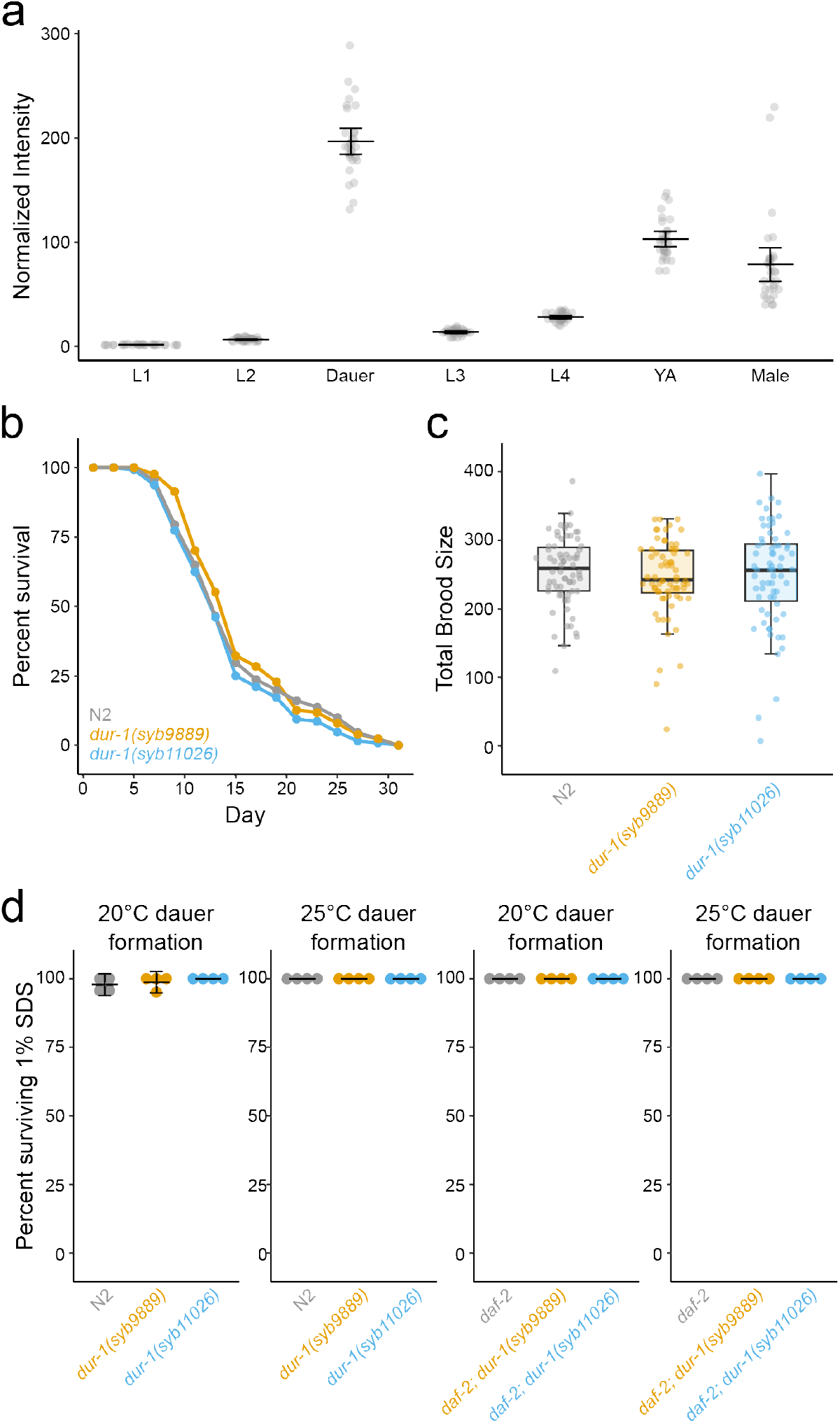
*dur-1* is primarily expressed during the dauer larval stage and does not disrupt normal physiology of *C. elegans*. a) Stage-specific quantification of *dur-1*::mNG translational fusion worms shows a significant increase in expression in dauer worms relative to other larval stages. Error bars depict 95% confidence intervals. b) Mutations in *dur-1* do not affect lifespan. Aggregate data is shown from four independent replicates. c) Mutations in *dur-1* do not impact brood size. d) *dur-1* mutants form SDS-resistant dauer larvae in wild-type (N2) and *daf-2(e1370)* backgrounds at both 20°C and 25°C.

We hypothesized that functions of DUR-1 may be unique to dauer larvae. However, since DUR-1 is also present in adult worms, we tested for other non-dauer phenotypes. Initially, we obtained an existing allele of *dur-1, ok1010* (strain VC687), that was reported to be highly sensitive to desiccation, and we backcrossed it five times to reduce background mutations [20]. Surprisingly, we isolated two lineages from the 5x backcrossed *dur-1(ok1010)* lines (designated isolate 1 and isolate 2). One of these was short-lived, had a reduced brood, and did not form SDS-resistant dauer larvae (Figure S2). These phenotypes were consistent with what was observed in the strain before any backcrossing. These worms had a propensity to hold their embryos and die as a “bag of worms” [31]. Multiple rounds of backcrossing and propagation either allowed for outcrossing of a linked background mutation or the isolation of a suppressor. To sidestep the ambiguity of these genetics, for further experiments we instead utilized a null mutant, *dur-1(syb9889)*, and a CRISPR-generated strain with the same *ok1010* deletion – *dur-1(syb11026)* (Figure S1).

We did not observe significant differences in lifespan of *dur-1(syb9889)* or *dur-1(syb11026)* worms compared to wild-type N2 worms (adjusted p-value N2 vs. *dur-1(syb9889)*=0.98, adjusted p-value N2 vs. *dur-1(syb11026)*=0.49, log-rank test, Figure 1b). There were also no differences in brood size of *dur-1* mutants and wild-type worms (p=0.92, one-way ANOVA, Figure 1c). These data suggest that *dur-1* does not contribute significantly to organismal physiology under standard developmental conditions.

Given the high levels of DUR-1 present in dauer larvae, we tested whether it is essential for dauer formation. Mutations in *lea-1*, the other LEA protein of *C. elegans*, led to temperature-sensitive defects in dauer formation, with worms grown at 25°C but not 20°C showing SDS sensitivity [29]. Therefore, we tested the null allele, *dur-1(syb9889)*, and partial deletion, *dur-1(syb11026)*, for an impact on dauer formation at both 20°C and 25°C. Using a standard SDS-resistance assay to monitor dauer formation, we observed near 100% survival of dauers exposed to 1% SDS for 30 minutes (Figure 1d) [32]. At both temperatures, dauer larvae did not show significant differences in SDS survival compared to wild-type N2 controls (Figure 1d). The same alleles of *dur-1* crossed into a *daf-2(e1370)* mutant to facilitate production of constitutive dauer larvae at 25°C similarly did not exhibit significant differences in survival relative to *daf-2(e3170)* controls (Figure 1d). We conclude that DUR-1 is not required for the development of SDS-resistant dauer larvae.

### DUR-1 expression increases during desiccation

Given the substantial increase in *dur-1* expression in dauer larvae, we were curious if there would be further upregulation in response to stress. We measured the fluorescence intensity of dauer larvae expressing *dur-1*::mNG and compared it with that of dauer larvae exposed to desiccation (4 days at 97.5% RH), osmotic stress (1 M NaCl), and heat stress (37°C). Desiccation resulted in an increase in DUR-1::mNG in both wild-type dauers formed by starvation at 20°C (p=0.01) and *daf-2(e1370)* mutant dauers formed constitutively by growth at 25°C (p<0.001, unpaired t-tests, Figure 2a,d). This finding agrees with a previous report that showed upregulation of *dur-1* during desiccation and suggests a concurrent increase in protein [20].

**Figure 2.**
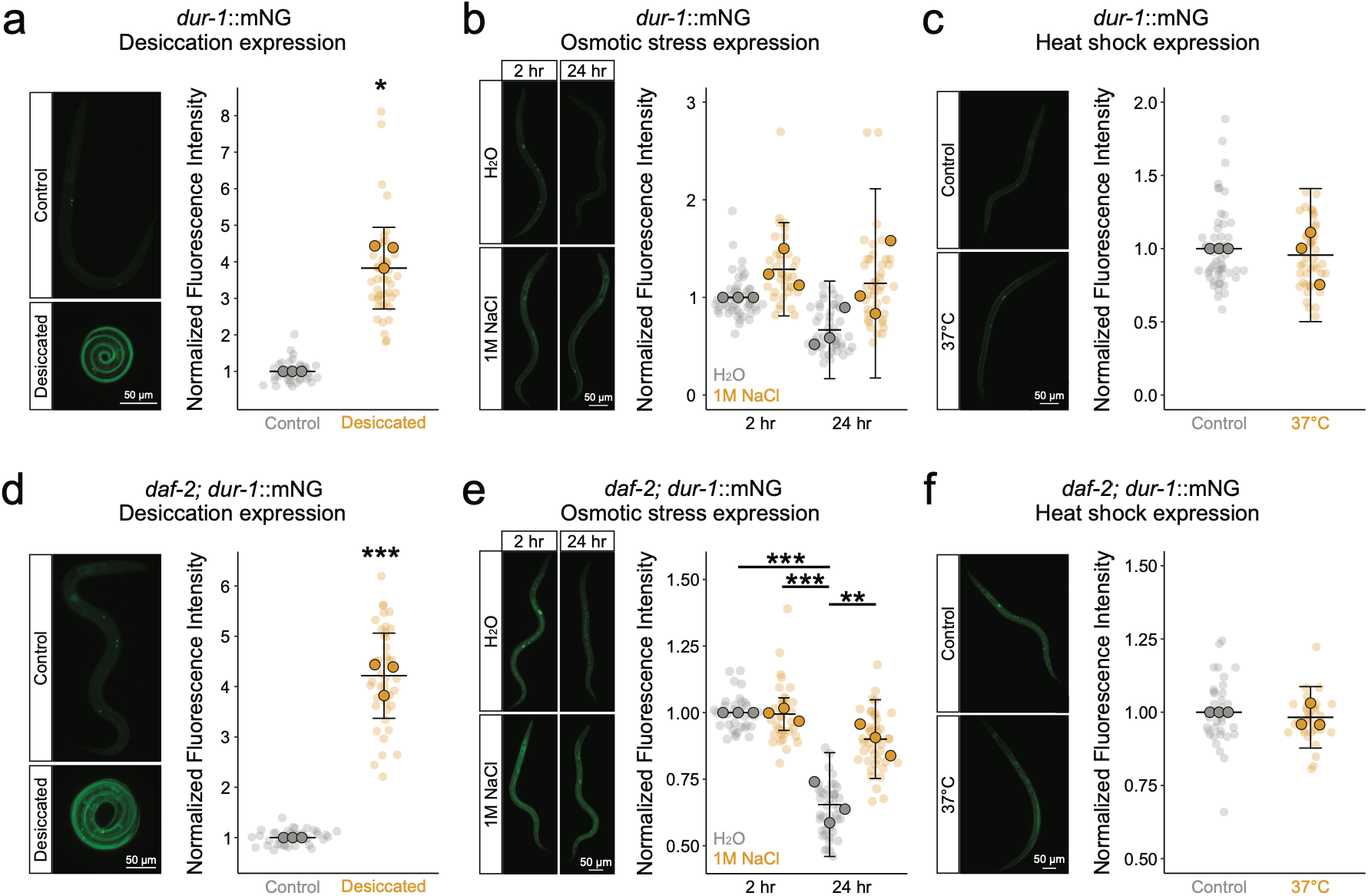
DUR-1 is upregulated by desiccation. a) Dauer worms from starved plates that were dried at 97.5%RH for 4 days show increased fluorescent intensity relative to control worms (p=0.01, unpaired T-test, n=3). b) There is no significant difference in expression during exposure to osmotic stress (1M NaCl) for 2 hr or for 24 hr. c) There is no change in fluorescent intensity (p=0.74, unpaired T-test, n=3) upon heat shock (6 hr at 37°C). d) Desiccated dauers that were formed constitutively in a *daf-2(e1370)* background at 25°C show increased *dur-1*::mNG fluorescence (p<0.001, unpaired T-test, n=3). e) *daf-2(e1370);dur-1*::mNG dauers formed at 25°C showed an apparent reduction in fluorescence intensity after 24 hr in water (p<0.001, Tukey’s Test, n=3), f) Worms in a *daf-2(e1370)* background also did not exhibit a significant change in DUR-1 expression following 6 hr at 37°C (p=0.59, unpaired T-test, n=3). * p<0.05, ** p<0.01, *** p<0.001.

Worms exposed to short-term osmotic stress (2 hr in 1M NaCl) did not have a significant change in DUR-1::mNG levels. After 24 hr, there was a reduction in DUR-1::mNG in control worms kept in water. In contrast, worms kept in 1M NaCl had less of a reduction in expression following 24 hr (Figure 2b,e). There were no significant changes in DUR-1 expression after 6 hr heat shock at 37°C in wild-type (p=0.74) or *daf-2(e1370)* dauers (p=0.59, unpaired t-tests, Figure 2c,f). These experiments support the conclusion that DUR-1, which is already upregulated in dauer larvae, is further induced by desiccation but not osmotic stress or heat shock.

### DUR-1 contributes to survival of desiccation and osmotic stress

Using the null mutant, *dur-1(syb9889)*, and putative hypomorph, *dur-1(syb11026),* we measured survival of dauer larvae exposed to three stresses – desiccation, osmotic stress, and heat shock. *C. elegans* dauer larvae can survive desiccation but require conditioning at relatively high humidity before exposure to lower humidities [19]. We measured desiccation survival after 4 days at 97.5% RH and after a subsequent day at 60% RH. We found that *dur-1(syb9889)*, but not *dur-1(syb11026)*, exhibited a reduction in desiccation survival at 97.5% RH in dauer larvae formed by starvation at 20°C (Figure 3a). Survival of worms, including N2 controls, was highly variable at 60%. The same *dur-1* mutants in combination with *daf-2(e1370)* showed a similar pattern, where *dur-1(syb9889)* but not *dur-1(syb11026)* was sensitive to desiccation – both at 97.5% RH and 60% RH (Figure 3a). We conclude that *dur-1* contributes to desiccation tolerance in dauers formed at 20°C.

**Figure 3.**
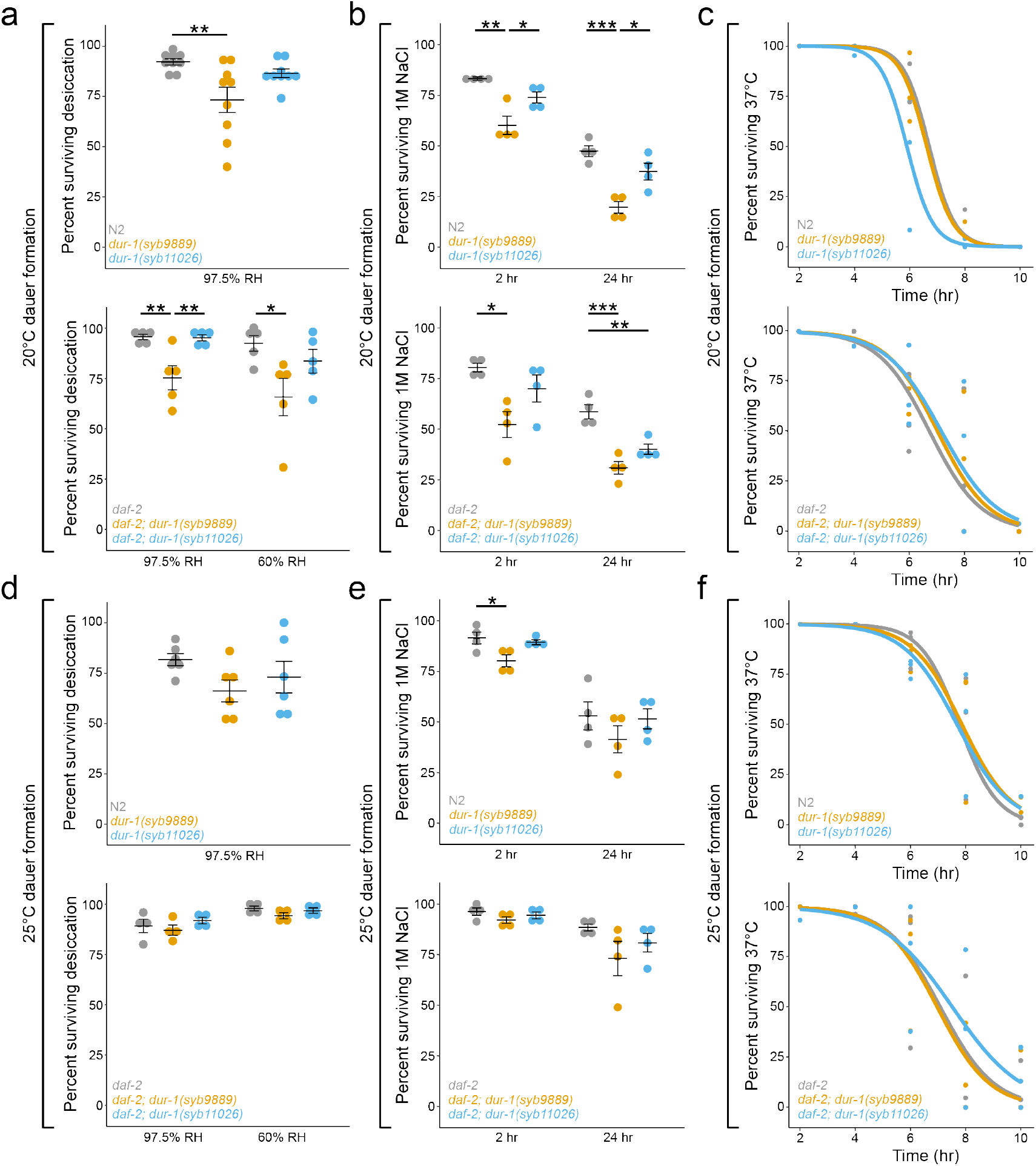
*dur-1* is required for dauer desiccation and osmotic stress survival at 20°C. a) *dur-1(syb9889)* mutants were sensitive to mild desiccation at 97.5% RH when dauer larvae were formed by starvation at 20°C (p=0.006, n=9). Similarly, in a *daf-2(e1370)* mutant background, *dur-1(syb9889)* dauers formed 20°C had reduced survival after drying at 97.5% RH and 60% RH. b) Compared to wild-type N2 worms, *dur-1(syb9889)* mutant dauer larvae formed at 20°C were sensitive to exposure to high salt (1M NaCl) for 2 hr (p=0.001, n=4) and 24 hr (p<0.001, n=4). In a *daf-2(e1370)* mutant background, *dur-1(syb9889)* dauers formed 20°C had reduced survival in 1M NaCl at 2 hr (p=0.01, n=4), and 24 hr (p<0.001, n=4). Additionally, *dur-1(syb11026)* mutants had reduced survival compared to N2 worms after 24 hr exposure to 1M NaCl (p=0.005, n=4). c) Mutations in *dur-1* had no impact on survival of dauers formed at 20°C that were exposed to heat shock at 37°C. d) There were no significant differences in desiccation survival (97.5% RH) between N2, *dur-1(syb9889)*, and *dur-1(syb9889)* mutants desiccation in dauer larvae formed by starvation at 25°C. *dur-1* mutant dauer larvae formed constitutively in a *daf-1(e1370)* background at 25°C did not show statistically distinguishable differences in desiccation survival at either 97.5% RH or 60% RH. e) *dur-1(syb9889)* mutant dauers formed at 25°C exhibited a modest survival defect relative to N2 when exposed to 1M NaCl for 2 hr (p=0.03, n=4). There were no significant differences in osmotic stress survival observed in *dur-1* mutants in combination with *daf-2(e1370)*. f) There were no significant differences in heat shock survival in dauer larvae formed at 25°C for either allele of *dur-1* in N2 or *daf-2(e1370)* genetic backgrounds. All p-values were calculated with Tukey’s post hoc tests following a significant 1-way ANOVA. * p<0.05, ** p<0.01, *** p<0.001.

Desiccation causes some similar subcellular stresses as osmotic stress and heat shock. For example, reduction of cellular volume and concentration of solutes are features of desiccation and osmotic stress, and challenges to proteostasis are common to all three stresses [13,33–37]. Dauer larvae were exposed to 1M NaCl for either 2hr or 24hr to determine the impact of *dur-1* on osmotic stress survival. After 2hr and 24hr, *dur-1(syb9889)* mutants had significantly lower survival than controls (Figure 3b). The same reduced osmotic stress survival of *dur-1(syb9889)* was observed in a *daf-2(e1370)* background. Additionally, *daf-2(e1370);dur-1(syb11026)* mutants had lower survival than *daf-2(e1370)* controls after 24 hr of exposure (Figure 3b). There was no difference in heat shock survival at 37°C in either *dur-1* allele relative to N2 or *daf-2(e1370)* controls (Figure 3c). These data indicate that *dur-1* contributes to desiccation and osmotic stress tolerance, but not heat shock survival in dauer larvae formed at 20°C.

The other LEA protein of *C. elegans*, LEA-1, has temperature-dependent effects on survival, with a dauer-defective phenotype at 25°C worsening survival of desiccation and osmotic stress[29]. We wondered if *dur-1* may have similar temperature-dependent effects on survival. Notably, we did not observe differences in dauer formation between temperatures (20°C vs. 25°C), nor in dauer formation mode (starvation vs. *daf-2* constitutive dauer) (Figure 1d). We repeated the same stress survival experiments in dauer larvae that were formed at 25°C. In this case, the *daf-2(e1370)* mutants form dauers constitutively, providing the second point of comparison. Surprisingly, we found that the impact of *dur-1* on dauer stress tolerance was blunted at 25°C relative to 20°C. *dur-1* mutant dauers formed at 25°C were not sensitive to desiccation (Figure 3d). *dur-1(syb9889)* mutant dauers only had a modest reduction in survival after 2 hr in 1M NaCl (p=0.03, Tukey’s test) and no statistical difference from controls after 24 hr (Figure 3e). In *daf-2(e1370)* constitutively formed dauers, there were no statistically distinguishable differences in osmotic stress survival. As was the case with *dur-1* mutant dauers formed at 20°C, there was no difference in heat shock survival in *dur-1* mutants formed at 25°C (Figure 3f). With the additional comparison of dauers formed at 25°C by starvation (N2 background) and constitutively (*daf-2* background), we conclude that temperature had a greater effect on survival than the mode of dauer formation. Altogether, these data support the model that *dur-1* promotes desiccation and osmotic stress tolerance of dauers formed at 20°C, but not 25°C.

Many desiccation-tolerant organisms have expanded families of LEA proteins. This is particularly true of plants, but even other invertebrates like tardigrades, rotifers, and brine shrimp have multiple LEA proteins [22,38]. *C. elegans* only has two known LEA-like proteins, providing a strategic opportunity to look at the interaction between LEA proteins *in vivo*. We created a *lea-1;dur-1* double mutant to determine if stress sensitivity is exacerbated beyond the effects seen in either single mutant. Because *lea-1* mutants do not form SDS-tolerant dauers at 25°C, and *dur-1* is uniquely functional at 20°C, we focused our analysis only on dauers formed at 20°C. Dauer larvae formed at 20°C did not have a dauer formation defect, even in *dur-1(syb9889);lea-1(cp423)* double mutants (Figure S3). As expected, *lea-1(cp423)* null mutants were sensitive to desiccation (Figure 4a). The *dur-1(syb9889);lea-1(cp423)* double mutant had significantly lower desiccation survival (8% on average) than each mutant individually (*dur-1(syb9889)*=77%, *lea-1(cp423)*=31%). Although the double mutant had lower desiccation survival than each single mutant, we lack the dynamic range to parse an additive versus synergistic effect of these mutations. In a *daf-2(e3170)* background the *dur-1(syb9889);lea-1(cp423)* double mutant combination worsened desiccation survival compared to *dur-1(syb9889)* alone but was not statistically distinguishable from *lea-1(cp423)*.

**Figure 4.**
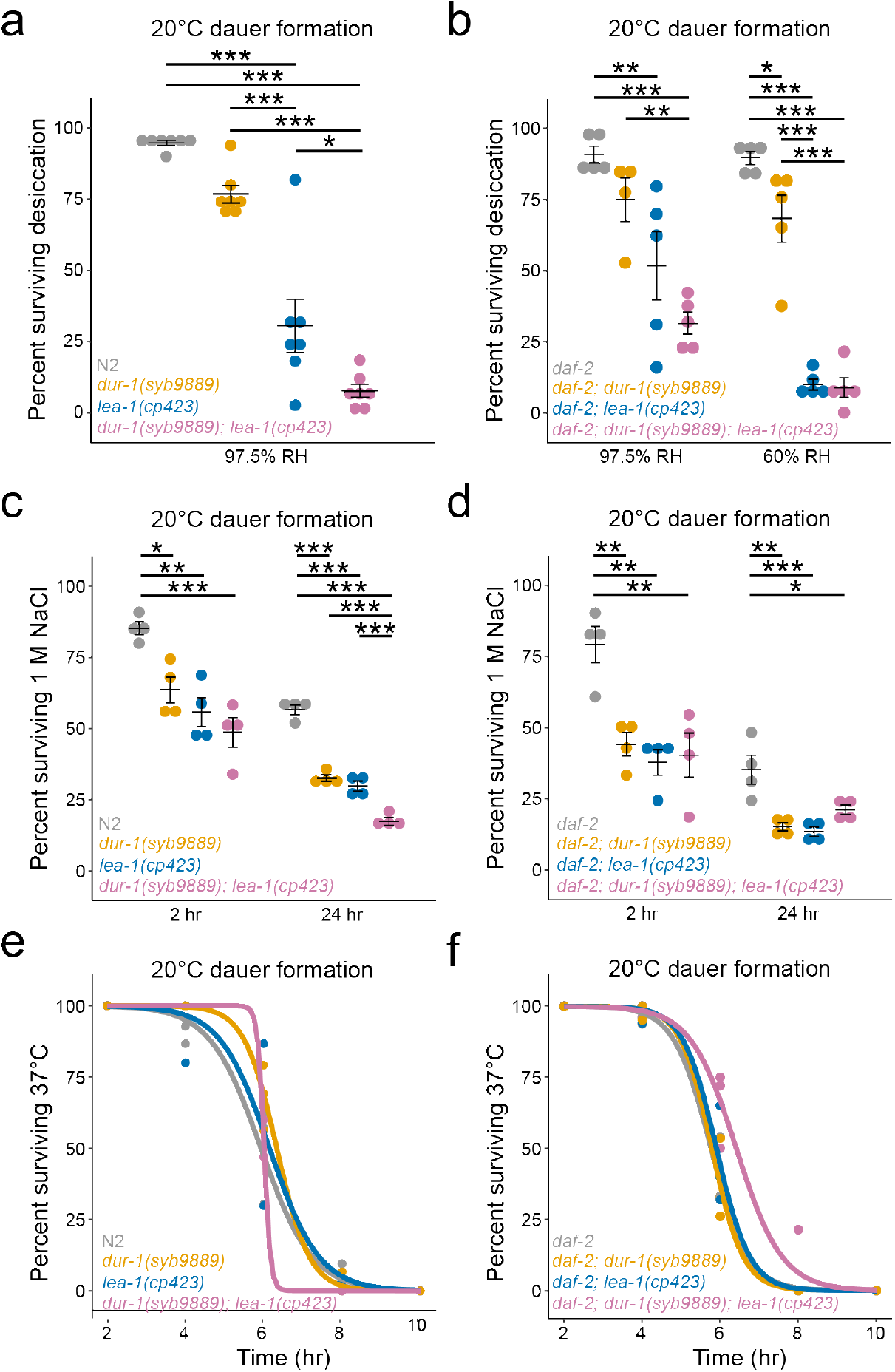
Combined *lea-1* and *dur-1* mutations increase stress sensitivity. a) After mild desiccation at 97.5% RH, *dur-1(syb9889);lea-1(cp423)* double mutants had reduced survival compared to N2 (p<0.001, n=7), *dur-1(syb9889)* single mutants (p<0.001, n=7), and *lea-1(cp423)* single mutants (p<0.001, n=7). b) In a *daf-2(e1370)* background, *dur-1(syb9889);lea-1(cp423)* mutants had reduced survival compared to *daf-2(e1370)* controls (p<0.001, n=5) and *dur-1(syb9889)* at both 97.5% RH and 60% RH. Survival of the *dur-1(syb9889);lea-1(cp423)* mutant was not detectably different from single *lea-1(cp423)* mutants. c) Exposure of *dur-1(syb9889);lea-1(cp423)* mutants to 1M NaCl for 2hr or 24hr resulted in reduced survival relative to N2 controls (p<0.001, n=4 in each case). Only after the 24hr exposure was survival of double mutants different from single *dur-1(syb9889)* (p<0.001, n=4) d) The same mutants in a *daf-2(e1370)* background had reduced osmotic stress survival relative to control animals following 2hr and 24hr of exposure to 1M NaCl. Survival of *dur-1(syb9889);lea-1(cp423)* double mutants and *dur-1(syb9889)* and *lea-1(cp423)* single mutant was statistically indistinguishable. e) Survival of 37°C heat shock is shown for N2 as well as *dur-1(syb9889)* and *lea-1(cp423)* single and double mutants. There was no significant difference in survival across genotypes. f) *dur-1(syb9889)* and *lea-1(cp423)* single and double mutants in a *daf-2(e1370)* background showed no significant differences in survival at 37°C. All p-values were calculated with Tukey’s post hoc tests following a significant 1-way ANOVA. * p<0.05, ** p<0.01, *** p<0.001.

Single mutants of *dur-1* and *lea-1* were sensitive to osmotic stress, as previously observed (Figure 3) [29]. The double mutant exhibited survival on par with the single mutants, with only longer exposure (24 hr) to 1M NaCl revealing a significant difference in survival between *dur-1(syb9889);lea-1(cp423)* mutants and each of the single mutants (Figure 4c). In combination with *daf-2(e1370)*, no differences were observed between single and double mutants. There was no significant difference in heat shock survival detected across genotypes in either an N2 or *daf-2(e1370)* genetic background (Figure 4e,f). In aggregate, these data support individual contributions of both *dur-1* and *lea-1* to desiccation and osmotic stress survival but do not suggest a strong synergistic relationship between these two LEA proteins.

### DUR-1 is a secreted basement membrane protein

To better understand the mechanism by which DUR-1 contributes to desiccation and osmotic stress survival, we sought to understand where it localizes to provide clues about what it may protect. Using an endogenous fluorescently tagged reporter strain, we visualized DUR-1 protein localization in *daf-2(e1370)* dauer worms (Figure 5). Dauer larvae exhibited fluorescence throughout the worm (Figure 5a), with apparent localization to the membranes of the body wall muscle cells (Figure 5b), the basement membrane of the pharynx (Figure 5c), the gonad (Figure 5d), and the hypodermis (Figure 5e). The localization patterns observed in *daf-2(e1370)* constitutive dauers formed at 25°C were also seen in dauer worms without a *daf-2* mutation that formed from starvation at 20°C (Figure S4). The strongest fluorescence intensity was visualized in three pairs of cells throughout the body that appeared to be coelomocytes, cells that endocytose proteins and macromolecules from the pseudocoelom [39,40]. We confirmed the identity of these cells by injecting a coelomocyte-specific RFP reporter (coel::RFP). Indeed, the mNG signal colocalized with the coel::RFP, indicating the accumulation of DUR-1 protein in coelomocytes (Figure 5f).

**Figure 5.**
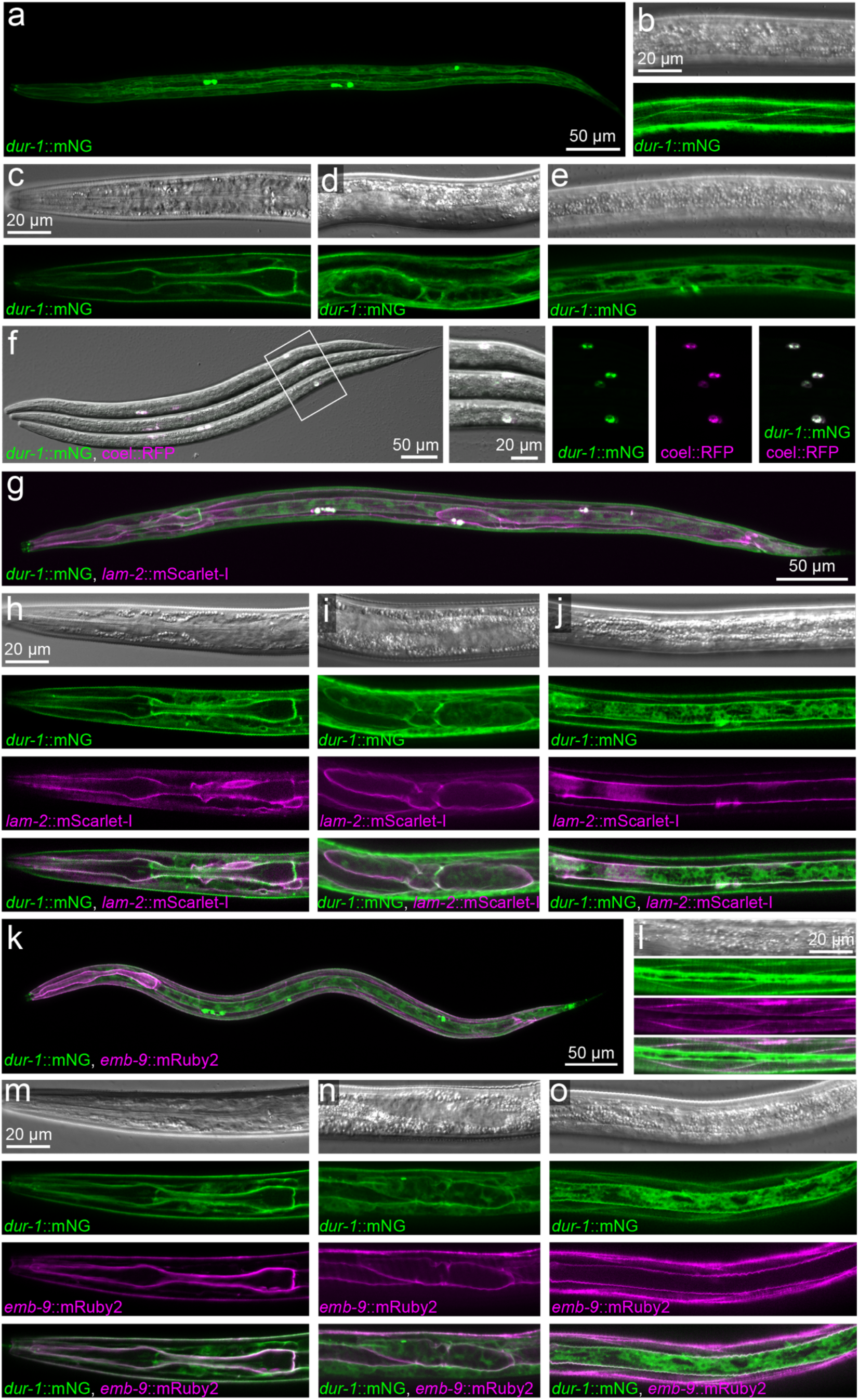
DUR-1 is a basement membrane protein. a) A representative image of a dauer larva with an endogenous fluorescent reporter (DUR-1::mNG). A maximum intensity projection shows fluorescence throughout the body. Fluorescence was visualized on the membranes of body wall muscle cells (b), the basement membrane surrounding the pharynx (b), the gonad (c), and the hypodermis (d). The scale bar in c applies to c-e. f) The strongest fluorescence signal was in the coelomocytes and demonstrated strong co-localization with coelomocyte-specific RFP. g) DUR-1 co-localizes with laminin (*lam-2*::mScarlet-I). Strong fluorescence signal was observed for both reporters in coelomocytes (g), and along basement membranes surrounding the pharynx (h), gonad (i), and hypodermis (j). k) DUR-1 co-localizes with a type IV collagen reporter (*emb-9*::mRuby2). Fluorescence was seen in body wall muscle cells (l), as well as basement membrane of the pharynx (m), gonad (n), and hypodermis (o). The scale bar in m applies to m-o. Note, all images show *daf-2(e1370)* mutant dauers formed constitutively at 25°C.

The accumulation of DUR-1 at sites of basement membrane (BM) deposition was surprising, as this protein was not previously reported, nor expected to be a BM component. To establish DUR-1 as a bona fide BM protein, we examined co-localization with hallmark basement membrane proteins, laminin (*lam-2)* and type IV collagen (*emb-9*) [41–43]. Fluorescence from the *lam-2::*mScarlet-I reporter was seen in coelomocytes and on basement membranes throughout the worm (Figure 5g). Sites of *dur-1*::mNG accumulation on basement membranes of the pharynx (Figure 5h), gonad (Figure 5i), and hypodermis (Figure 5j) each showed overlap with laminin. Of note, *dur-1*::mNG was also present beyond the BM, particularly in the hypodermis (Figure 5j).

To further validate the BM localization of DUR-1, we examined co-localization with type IV collagen (*emb-9*). The *emb-9*::mRuby2 fluorescence pattern was highly similar to that of laminin but with less pronounced localization to the coelomocytes and more signal in the body wall muscle (Figure 5k,l). The fluorescence signal of *dur-1*::mNG co-localized with the *emb-9*::mRuby2 on BM of the pharynx (Figure 5m), gonad (Figure 5n), and hypodermis (Figure 5o). The colocalization of *dur-1*::mNG with two core BM proteins, laminin and type IV collagen, establishes DUR-1 as a component of the BM.

Basement membrane proteins are typically secreted proteins. Accumulation in the coelomocytes, as seen with DUR-1 (Figure 5f), is also a hallmark of secreted proteins. DUR-1 has a predicted secretion signal peptide, which further suggests that DUR-1 is secreted (Figure S5) [44]. To determine *dur-1* expression sites, we created a transcriptional reporter by fusing 1kb upstream of *dur-1* to mKate2 (*dur-1p*::mKate2). This reporter was introduced as a single-copy insertion on Chr I. Worms co-labeled with the transcriptional and translational reporters show distinct patterns of fluorescence (Figure 6). Expression of the *dur-1p*::mKate2 construct was restricted primarily to body wall muscle (Figure 6a), with no expression in the pharynx or other tissues where DUR-1::mNG was seen (Figure 6b). Within muscle cells, the mKate2 signal was cytosolic, in contrast to the membrane-localized DUR-1::mNG (Figure 6c). Collectively, these expression patterns reveal that *dur-1* is primarily produced in body wall muscle and is secreted throughout the body, accumulating in coelomocytes and on basement membranes.

**Figure 6.**
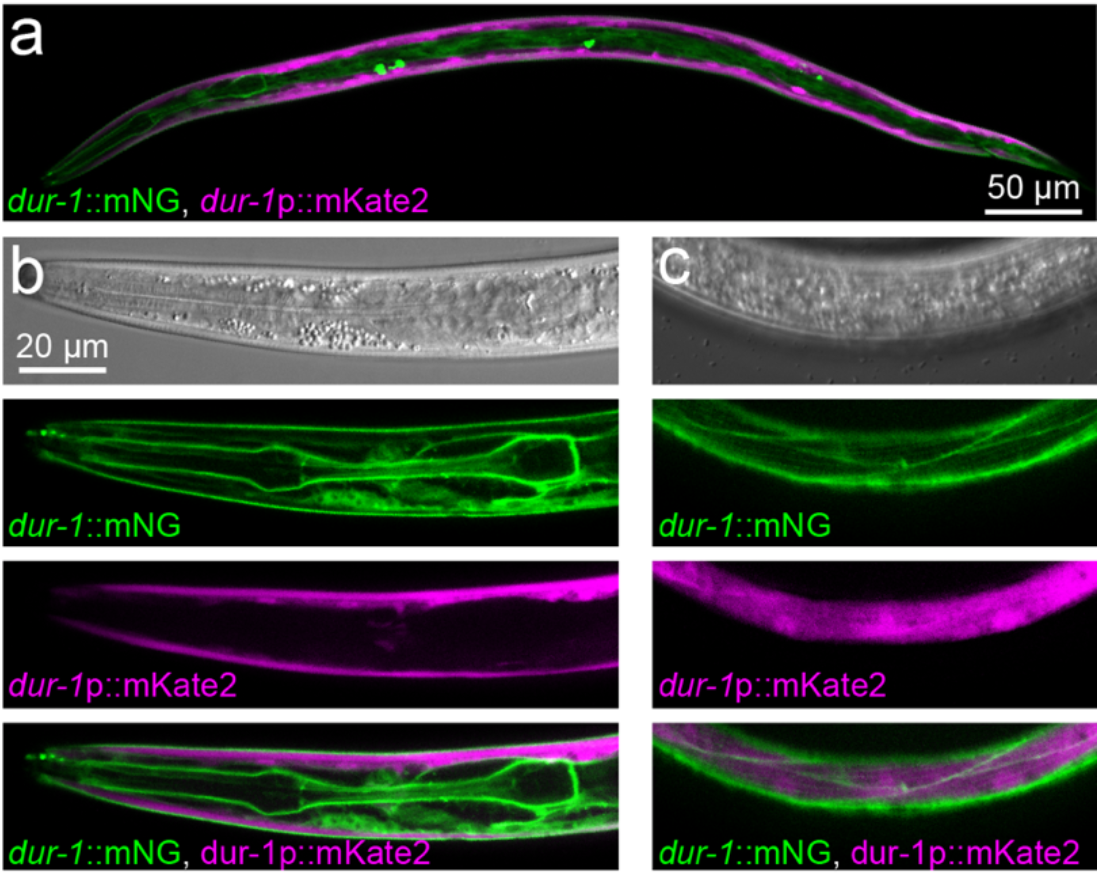
*dur-1* is transcribed in and secreted from the body wall muscle. a-b) A transcriptional reporter fusing 1kb of sequence upstream of *dur-1* with mKate2 reveals expression that does not overlap with the translational fusion. The transcriptional reporter is strongly expressed in the body wall muscle of dauer larvae. c) The transcriptional reporter is localized to the cytosol of muscle cells, whereas the translational fusion is present on the membranes of the same cells. The scale bar in b also applies to c.

### DUR-1 promotes tissue integrity during stress

The surprising localization of DUR-1 to basement membranes led us to hypothesize that it may reinforce basement membranes during mechanical stress associated with desiccation and osmotic stress. To test this hypothesis, we used the pharynx as a readily observable and deformable tissue. Visualization of pharynxes was enhanced by use of the type IV collagen (*emb-9::mRuby2)* reporter. Control animals that were not exposed to stress had pharyngeal morphology typical of dauer larvae and no baseline morphological changes in *dur-1(syb9889)* mutants (Figure 7a,e). Upon desiccation, most pharynxes in control (*daf-2)* and *dur-1(syb9889)* mutant animals contracted and exhibited severe deformation (Figure 7b). Due to the severity of the phenotype even in control animals, there was not a significant increase in deformity of pharynxes in *dur-1(syb9889)* mutants. Exposure to osmotic stress (1M NaCl) also resulted in a higher percentage of deformed pharynxes and significantly more deformities than in controls (Figure 7c,e). Heat shock did not impact the morphology of the pharynxes (Figure 7d,e).

**Figure 7.**
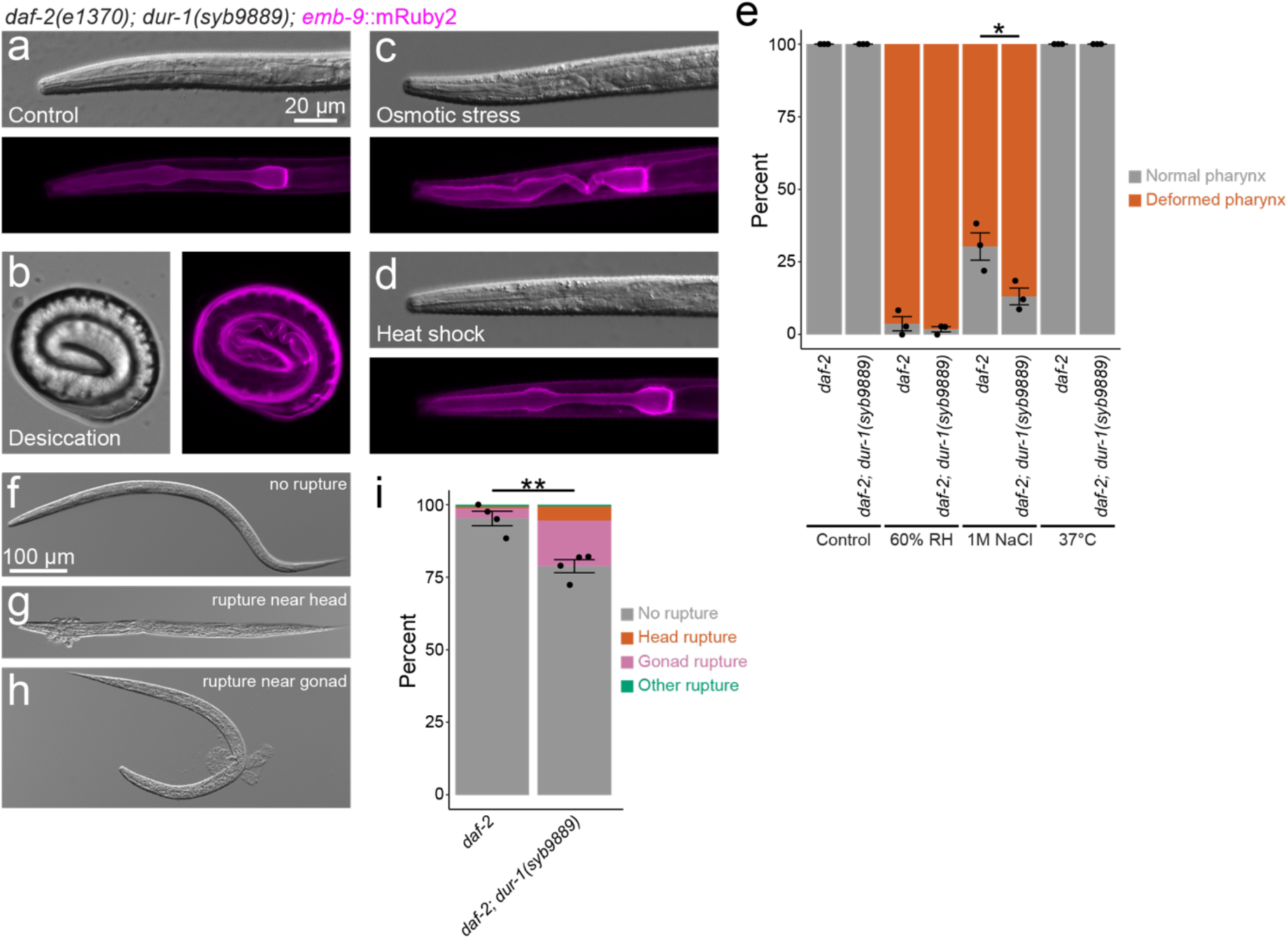
*dur-1* promotes tissue integrity during stress. The type IV collagen reporter (*emb-9*::mRuby2) facilitated visualization and scoring of pharynx phenotypes of *dur-1(syb9889)* mutants and *dur-1* WT worms, both in combination with *daf-2(e1370)*. a) A representative image shows a normal pharynx in a control animal. b) An animal following desiccation at 97.5% RH shows a severely deformed pharynx. c) A representative deformed pharynx after osmotic stress (1M NaCl for 2 hr), shows bending in the isthmus. d) Animals subjected to heat shock (37°C) retained normal pharynx morphology. e) Quantification of the percentage of normal and deformed pharynxes across control and stress conditions reveals a lower frequency of intact pharyngeal structure in *daf-2(e1370); dur-1(syb9889)* mutants following a 2 hr exposure to 1M NaCl (p=0.04, unpaired t-test, n=3). Following desiccation, most animals recovered with no rupture in the cuticle (f). Upon rehydration, some worms burst near the head (g) or gonad (h). i) The frequency of these phenotypes is quantified and shows a significant increase in the percentage of worms with rupture phenotypes in *dur-1(syb9889)* mutant animals (p=0.003, unpaired T-test, n=4). * p<0.05 ** p<0.01.

Cells can rupture upon desiccation and rehydration, and we reasoned that a similar phenomenon may occur at the level of tissues [45,46]. We observed instances of organs rupturing from worms upon rehydration and recovery from desiccation. We hypothesized that weakened basement membranes and limited extracellular protection may augment such phenotypes. In control animals, there were relatively few instances of rupture, with the most common site being around the gonad (Figure 7f-i). In *dur-1* mutant animals, there was a significant increase in the frequency of tissue rupture following 1 hr of rehydration (Fig. 7i). This result supports a model in which DUR-1 serves as an extracellular protectant that associates with basement membranes and promotes tissue integrity in support of stress survival.

## Discussion

Late embryogenesis abundant (LEA) proteins are known to be desiccation protectants. However, the mechanisms by which they function can vary. For example, they can both limit protein aggregation and stabilize cell membranes [15,26,27,47–52]. Here, we report a new mode of protection for the secreted LEA-like protein, DUR-1, which we show is a component of the basement membrane (BM) that stabilizes tissues. This finding establishes a role for a LEA protein that acts in a cell-nonautonomous manner and highlights the importance of extracellular protection during stress.

DUR-1 is significantly upregulated during the stress-tolerant dauer diapause stage of *C. elegans* (Figure 1a). The dauer-enriched expression profile of the endogenous *dur-1*::mNG reporter may be useful as a visual marker for isolating dauer larvae [53,54]. *dur-1(ok1010)* mutants were previously reported to be strongly desiccation-sensitive [20]. We found that clean genetic alleles are sensitive to desiccation, but not to the same degree as reported (Figure S2). We found that DUR-1 is required for osmotic stress survival but is dispensable for heat shock survival (Figure 3). Interestingly, temperature also contributed to survival phenotypes, with dauer larvae formed at 20°C being more sensitive than those formed at 25°C (Figure 3). It is conceivable that at higher temperatures, hormetic mechanisms, such as upregulation of chaperones or oxidative stress response proteins, may prime worms for enhanced desiccation and osmotic stress survival [55]. Other protectants induced by elevated temperature (25°C) may mask a role of DUR-1 in supporting survival. Testing this model by measuring transcriptional or translational changes between dauers formed at each temperature may allow for the definition of new protectants that could act redundantly with DUR-1.

We observed stronger desiccation and osmotic stress-sensitive phenotypes in *dur-1(syb9889)* null mutants than the hypomorphic *ok1010/syb11026* alleles (Figure 3). The DUR-1 protein contains an N-terminal signal peptide, a long stretch of disorder that includes a predicted prion-like domain, and several repeated 22-mer motifs (Figure S6). Such motifs are typical of LEA-like proteins [29,56]. Because the *syb11026* allele does not delete the prion-like domain, nor completely ablate the stretch of 22-mers, it is not clear if those regions of the protein are essential for DUR-1 function. Given the differences in survival between the null and the hypomorph, these regions may be important for eliciting protection and/or establishing the localization of the protein.

LEA proteins can be found in diverse cellular compartments and organelles in plants and animals [57–62]. Most LEA proteins remain within cells, but some contain predicted signal peptides for secretion and, in some cases, have been shown to be secreted [57,58,63]. Although there is precedent for secretion of some LEA-like proteins, localization and functions outside the cell remain poorly characterized [57,58,64]. Our ability to visualize endogenous DUR-1 *in vivo* allowed us to determine that it is secreted (as seen in the discrepancy between transcriptional and translational reporters, Figure 6) and localizes to the basement membrane. DUR-1 is the first example of a LEA-like protein associated with the BM. This raises the possibility that other secreted LEA proteins may also interact with BM or other components of the extracellular matrix.

Secreted proteins beyond the LEA family can also contribute to extracellular protection. Tardigrade abundant heat soluble proteins are another class of natively disordered proteins that, similarly, have diversified in compartmentalization between the cytosol (CAHS), mitochondria (MAHS), and secreted versions (SAHS) [65–70]. The SAHS proteins are more structured than CAHS and MAHS proteins, exhibiting structural similarity to fatty acid binding proteins [71,72]. SAHS proteins can protect against fusion of dried liposomes and contribute to bacterial desiccation tolerance [73]. It is not known if or how SAHS proteins or other secreted protectants may interact with the extracellular matrix. Nonetheless, this parallel example of spatial diversification of protectant localization reinforces the notion that understudied extracellular protectants likely play important roles in supporting animal stress tolerance.

The stress sensitivities observed in *dur-1* mutants suggest a model in which DUR-1 mitigates damage common to osmotic stress and desiccation but distinct from heat stress (Figure 3). During osmotic stress and desiccation, water leaves cells, causing contraction and mechanical stress on tissues composed of, or in contact with, these shrinking cells [74,75]. As one conspicuous example, we observed deformation of the pharynx upon desiccation and osmotic stress (Figure 7). The localization of DUR-1 to basement membranes, including BM surrounding the pharynx, suggests that it could stabilize tissues during these stresses. Therefore, we hypothesize that DUR-1 serves to reinforce tissue integrity under mechanical stress. Indeed, loss of *dur-1* led to reduced integrity of the pharynx in worms exposed to high salt. Furthermore, the loss of *dur-1* led to an increased frequency of cells and tissues bursting out of the worms upon rehydration and recovery from desiccation. We suspect that this may be due in part to weakened basement membranes and perhaps differences in cuticle integrity. Future work to better understand how DUR-1 integrates into the basement membrane and how it affects tensile strength, stiffness, elasticity, and other properties will be important for a deeper understanding of BM remodeling during stress [8,76,77].

## Materials and Methods

### C. elegans maintenance

Worms were maintained according to standard laboratory conditions on nematode growth media (NGM) plates and fed OP50 *E. coli* [78]. Strains in the N2 genetic background were maintained at 20°C and 25°C, and strains in the *daf-2* genetic background were maintained at 15°C and 20°C. Males were generated by putting L4 worms at 32°C for 5–6 hrs and 25°C overnight, and standard methods were employed for crossing worms. The following strains and alleles were used: N2, CB1370 *daf-2(e1370)*, PHX9842 *dur-1*(*syb9842*[*dur-1*::mNG]) IV, HLS1 *dur-1(ok1010)* IV (5x backcross sick line), HLS2, *dur-1(ok1010)* IV (5x backcross healthy line), HLS3 *daf-2(e1370)* III*; dur-1(syb9889)* IV, HLS4 *daf-2(e1370)* III*; dur-1(ok1010)* IV (5x backcross sick line), HLS5 *daf-2(e1370)* III*; dur-1(ok1010)* IV (5x backcross healthy line), HLS6 *daf-2(e1370)* III*; dur-1(syb9842[dur-1::mNG])* IV, HLS13 *dur-1(syb9889)* IV (1x backcross of this allele), HLS22 *dur-1(syb9889)* IV*; lea-1ko(cp423[Pmyo-2>GFP-C1::myo-2 3’ UTR lox511I])* V, HLS23 *dur-1(syb11026)* IV (1x backcross), HLS24 *daf-2(e1370)* III; *dur-1(syb11026)* IV, LP847 *lea-1ko*(cp423[Pmyo-2>GFP-C1::myo-2 3’ UTR lox511I]) V, LP852 *daf-2(e1370)* III; lea-1ko(cp423[Pmyo-2>GFP-C1::myo-2 3’ UTR lox511I]) V, HLS8 *daf-2(e1370)* III*; dur-1(syb9889)* IV*; lea-1ko(cp423[Pmyo-2>GFP-C1::myo-2 3’ UTR lox511I])* V, HLS12 *hlaSi1[dur-1p::mkate2::tbb-2 3’UTR] I; daf-2(e1370)* III*; dur-1(syb9842[dur-1::mNG])* IV, HLS28 *daf-2(e1370)* III; *dur-1(syb9842*[*dur-1*::mNG]) IV; *lam-2(qy154*[*lam-2*::mScarlet-I]) X, HLS30 *daf-2(e1370)* III; *qy244*[*emb-9*::mRuby2] III, HLS31 *daf-2(e1370)* III; qy244[emb-9::mRuby2] III; dur-1(syb9842[dur-1::mNG]) IV, HLS32 *daf-2(e1370)* III; qy244[*emb-9*::mRuby2] III; dur-1(syb9889) IV.

### Formation of dauer larvae and SDS resistance

For all strains at 20°C, dauers were picked from starved plates maintained at 20°C. For N2 background strains at 25°C, dauers were picked from starved plates maintained at 25°C. For *daf-2(e1370)* background strains at 25°C, embryos were picked from plates maintained at 15°C and shifted to 25°C for 5 days. To test SDS resistance of dauer larvae, ~50 dauer worms per genotype were picked into 1 mL of 1% SDS in a 24-well plate. After 30 min, worms were scored for viability by movement in response to gentle prodding.

### Lifespan

For lifespan assays, embryos were picked onto fresh NGM plates with OP50 *E. coli*. For each genotype, 6–7 plates of five embryos each were included to start each replicate. Worms were maintained at 20°C and transferred to new NGM plates every other day until they were no longer reproductive. Worms that crawled onto the sides of plates or were otherwise missing were censored. Survival was scored every other day by gently tapping worms and assessing movement. Four independent biological replicates were conducted.

### Desiccation, osmotic stress, and heat shock

Worms were desiccated according to a previously established protocol [19,29]. Desiccation chambers were created using varying ratios of glycerol to water to produce 97.5% and 60% relative humidity (RH). Desiccation chambers were kept at either 20°C or 25°C for the duration of experiments to match the temperature at which dauers were formed. Dauer worms were picked into 1.5 µL droplets of water and exposed to 97.5% RH for 4 days as preconditioning. Worms were then either rehydrated in water or moved to 60% RH chambers for one day before rehydration. Worms were allowed to recover for at least 4 hrs before being scored for viability by movement in response to physical stimulus.

To test osmotic stress survival, dauer worms were picked into 1 mL of 1M NaCl in a 24-well plate. Dauer larvae were also picked into 1 mL of water as a control for the impact of time spent in liquid culture. Worms were scored for viability based on movement in response to physical stimulus after 2 hrs and 24 hrs.

To measure heat shock survival, dauer worms were picked onto multiple plates and shifted to 37°C. Plates were removed from 37°C incubator at 2, 4, 6, 8 and 10 hr intervals and scored for viability based on movement in response to gentle prodding after 1 day of recovery.

### Fluorescent imaging and quantification

To visualize fluorescent reporters, worms were imaged with a Zeiss AxioObserver outfitted with a Viluma7 light source, Apotome 3, and Axiocam 820 mono camera. Images were taken with a 10x or 20x Plan-Apochromat objective or C-Apochromat 40x objective with a water auto-immersion module.

For fluorescence quantification, worms were imaged with a 10x or 20x objective. Images were imported into FIJI for analysis. Whole worms were outlined manually, and total fluorescence intensity was measured. The outline of the worm was moved to a background area of the image, and the total fluorescence intensity was measured again to obtain a background measurement. The fluorescence intensity was calculated by subtracting the background value from that of the worm. The relative fold change in fluorescence was calculated by dividing individual worm measurements by the average of the control samples for that replicate.

### Statistical analysis

Statistical tests used for each analysis are noted in figure legends. Unpaired t-tests were conducted in Microsoft Excel. ANOVAs and post hoc Tukey’s tests were conducted in RStudio. Lifespan analysis was conducted on aggregate data from four biological replicates with OASIS 2, and data were plotted with RStudio [79,80].

## Data Availability

The data that support the findings of this study are available in supplemental data file 1.

## Acknowledgements

We thank Capri Woss for assistance in starting the project. We thank Dave Sherwood for helpful discussions and feedback and the Sherwood Lab at Duke University for sharing strains NK3072 and NK2767. Alleles *syb9889*, *syb9842*, and *syb11026* were created by SunyBiotech. Some strains were provided by the CGC, which is funded by NIH Office of Research Infrastructure Programs (P40 OD010440).

## Author contributions

BAB: Conceptualization, Methodology, Formal analysis, Investigation, Data curation, Writing – review & editing, Visualization. ALL: Investigation, Writing – review & editing. MKL: Investigation, Writing – review & editing. ZW: Investigation, Writing – review & editing. JDH: Conceptualization, Formal analysis, Investigation, Data curation, Writing – original draft, Writing – review & editing, Visualization, Supervision, Project administration, Funding acquisition.

## Figures and Figure Legends

**Figure S1.**
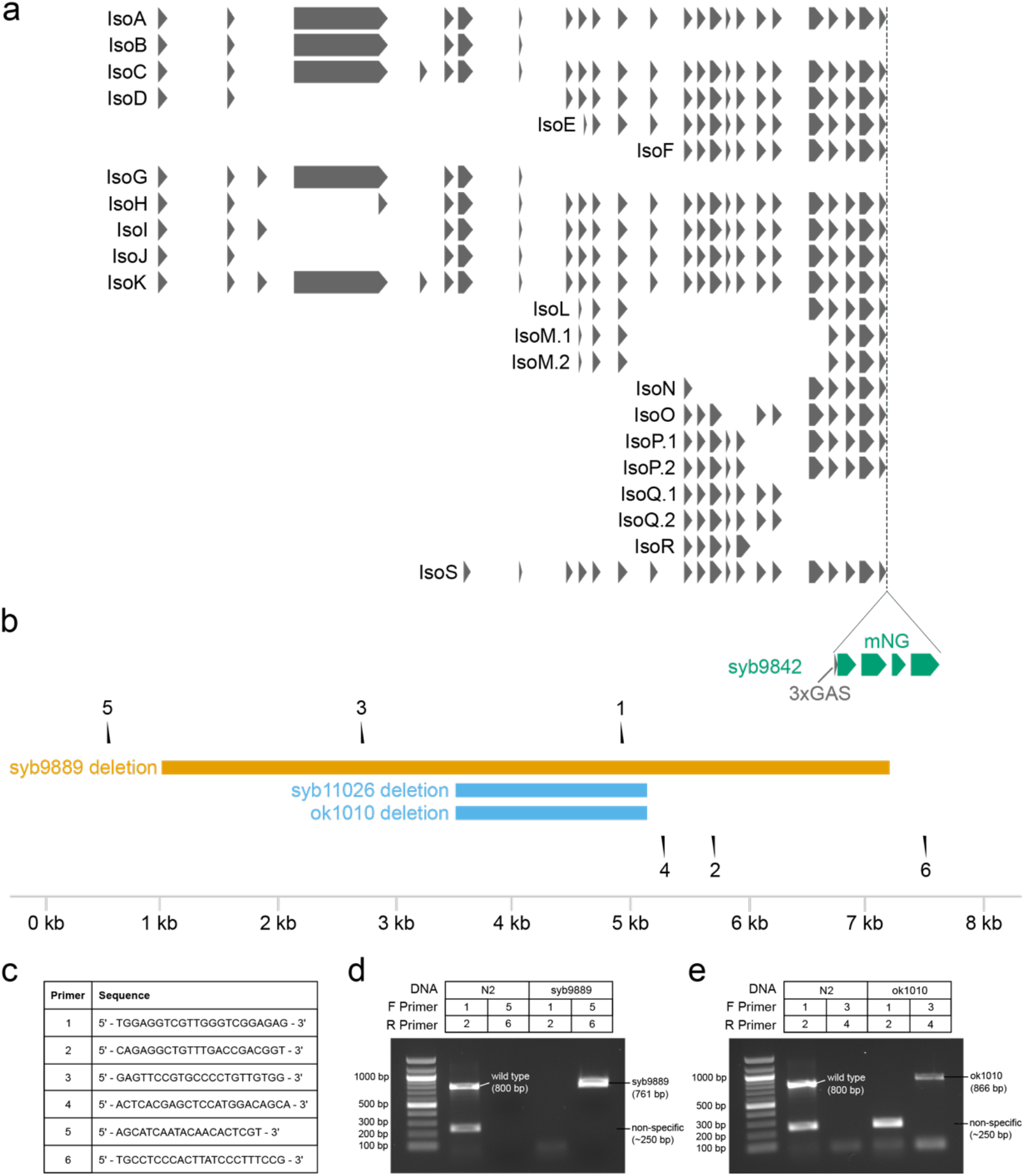
A schematic of the gene structure and mutant alleles of *dur-1*. a) *dur-1* has many predicted isoforms. b) An existing mutant allele, *dur-1(ok1010)*, was obtained, and a new null allele *dur-1(syb9889)* and an allele identical to *ok1010, dur-1(syb11026)*, were obtained. Additionally, the insertion of fluorescent mNG at the C-terminus of the protein, *dur-1(syb9842)*, allows for visualization of the endogenous protein. Locations of primers for genotyping each of these genomic alterations are indicated. c) Primers for genotyping alleles of *dur-1*. d) An example gel shows the genotyping strategy for *dur-1(syb9889)*. e) Genotyping strategy for *dur-1(ok1010)* and *dur-1(syb11026)*.

**Figure S2.**
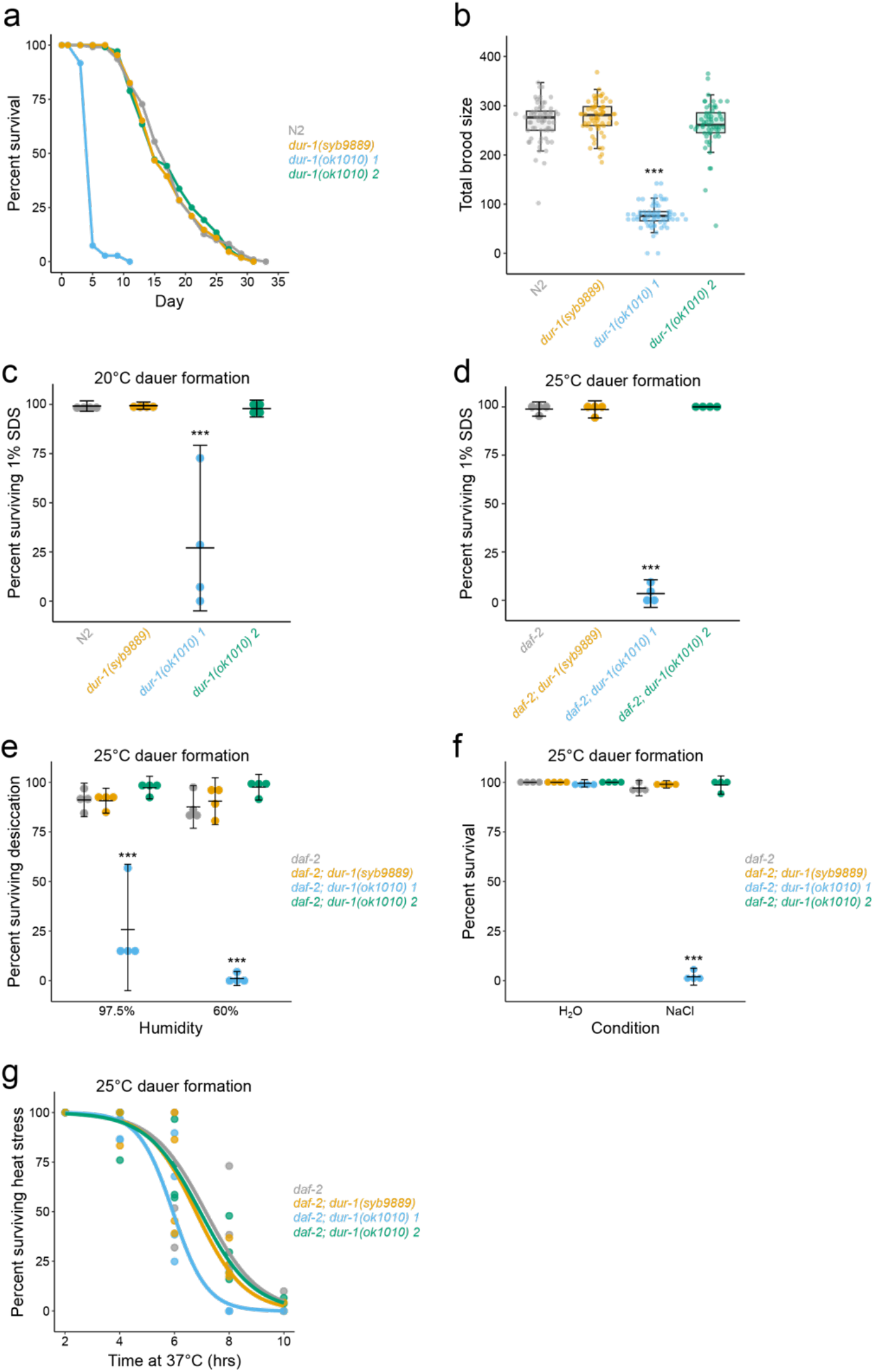
Following five rounds of back-crossing to N2, two *dur-1(ok1010)* isolates show discrepant phenotypes. a) The lifespan of each *dur-1(ok1010)* isolate is shown alongside *dur-1(syb9889)* and WT (N2) worms. Isolate 1, but not 2 of *dur-1(ok1010)* was short-lived (p<0.001, Log-Rank Test), with a mean lifespan of 5 days. b) Brood size of the same strains in A is plotted. Isolate 1 of *dur-1(ok1010)* had significantly reduced brood (p<0.001, Tukey’s Test). c) SDS survival of dauer worms for each *dur-1* mutant/isolate suggest a deficiency in dauer formation of *dur-1(ok1010)* isolate 1 on starved plates at 20 °C. d) SDS survival of dauer worms for each *dur-1* mutant/isolate in a *daf-2(e1370)* background reveals a defect in dauer formation in *dur-1(ok1010)* isolate 1 at 25 °C. e) Desiccation survival of *daf-2(e1370)* background dauer worms shows compromised survival of *dur-1(ok1010)* isolate 1 (p<0.001, Tukey’s Test, n=4). Desiccation survival of *dur-1(ok1010)* isolate 2 was statistically indistinguishable from *daf-2(e1370)* controls. f) Osmotic stress survival (2 hr, 1M NaCl) of *daf-2* background dauer worms shows sensitivity of *dur-1(ok1010)* isolate 1 (p<0.001, Tukey’s Test, n=4), but not isolate 2. g) Survival over time is plotted for *daf-2(e1370)* background dauer worms exposed to 37 °C heat shock. Curves were fit to individual data points from four experiments. There was no significant effect of genotype on survival (p=0.13, 1-way ANOVA, n=4).

**Figure S3.**
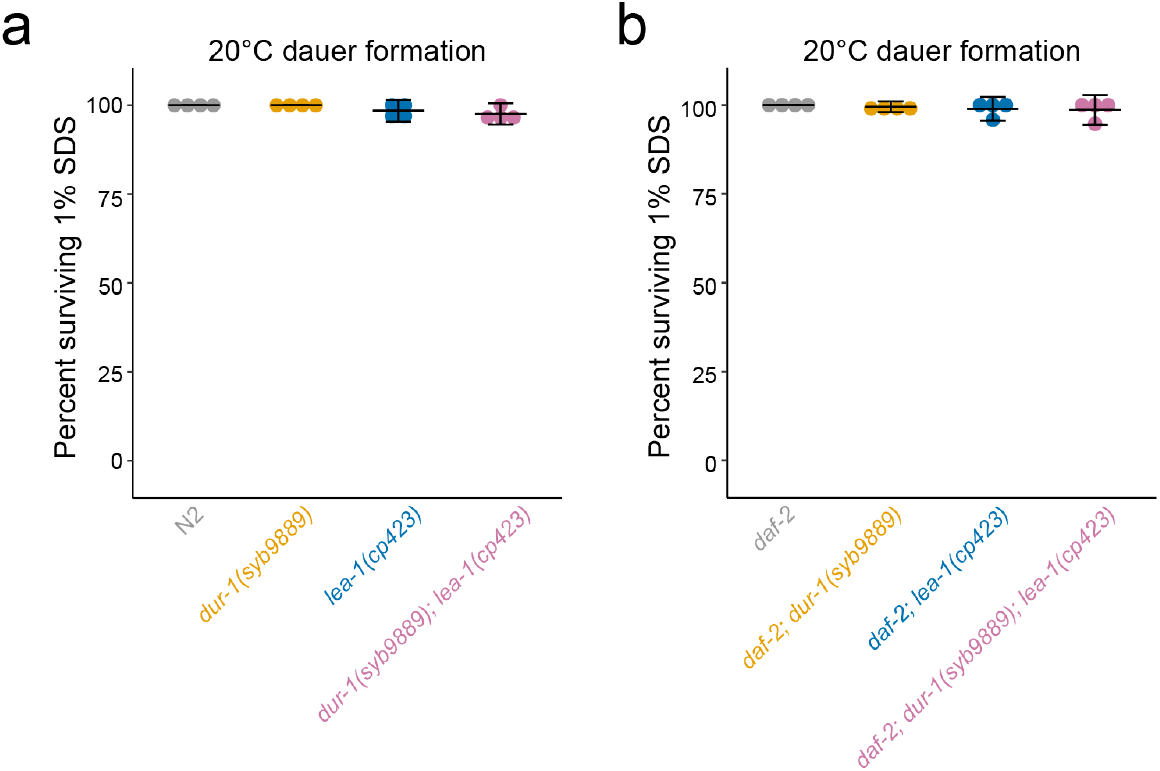
Combined mutations in *dur-1* and *lea-1* do not disrupt dauer formation at 20°C. a) In a wild-type (N2) background, *dur-1* and *lea-1* mutations individually and in combination do not sensitize dauer worms to SDS. b) In a *daf-2(e1370)* background, *dur-1* and *lea-1* individual and combination mutants form SDS-resistant dauer larvae at 20°C.

**Figure S4.**
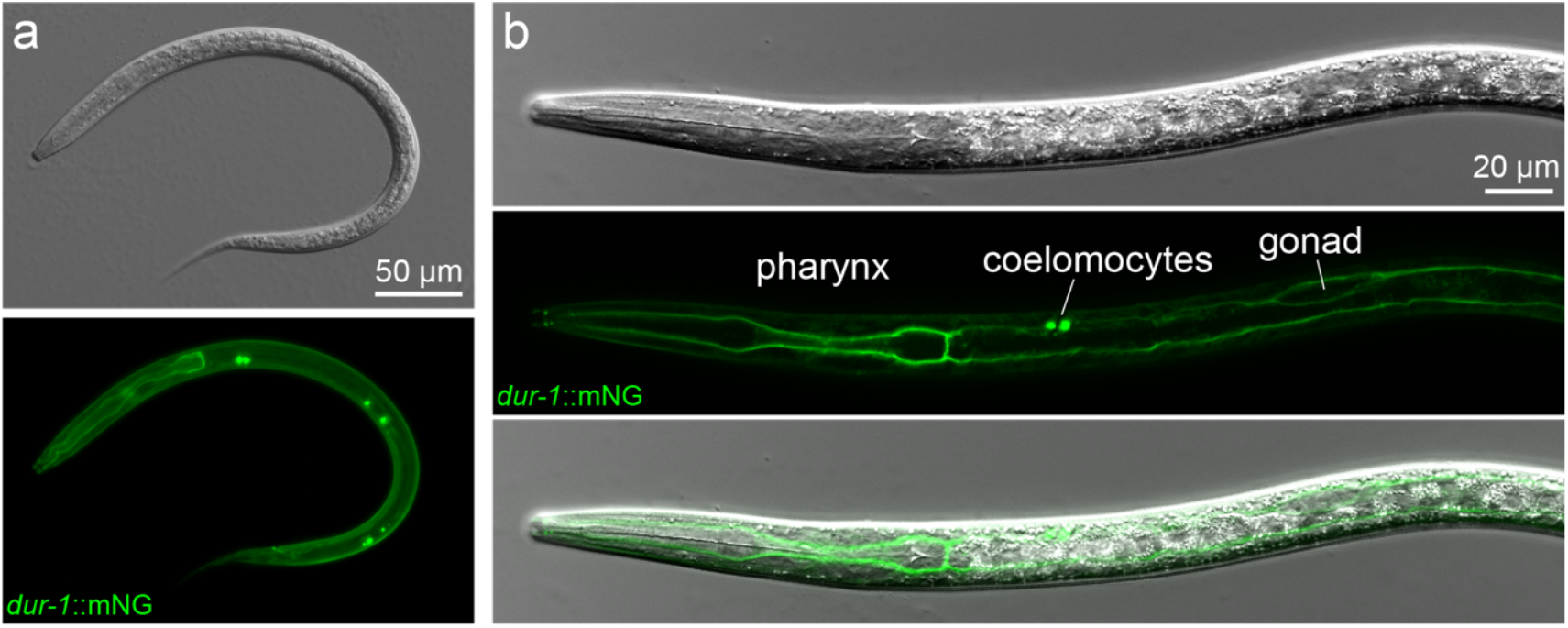
DUR-1::mNG expression and localization is consistent at 20 °C in a WT background. a) A maximum intensity projection of a dauer worm formed by starvation at 20 °C shows DUR-1-tagged protein localized to coelomocytes, the basement membrane of the pharynx, and throughout the body cavity. b) A higher-magnification image depicts the localization of DUR-1 to the basement membrane surrounding the pharynx, intestine, and gonad, in addition to coelomocytes.

**Figure S5.**
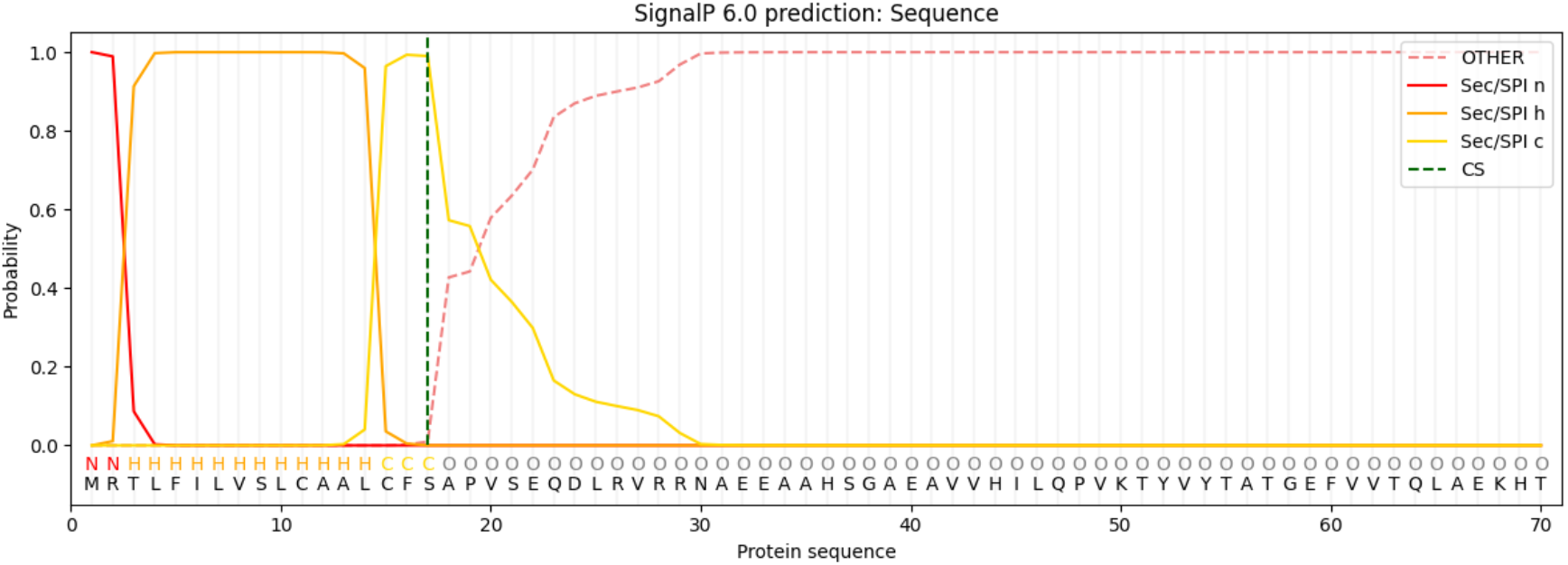
A predicted signal peptide suggests that DUR-1 is a secreted protein. Isoform K of DUR-1 was used for the prediction in SignalP 6.0.

**Figure S6.**
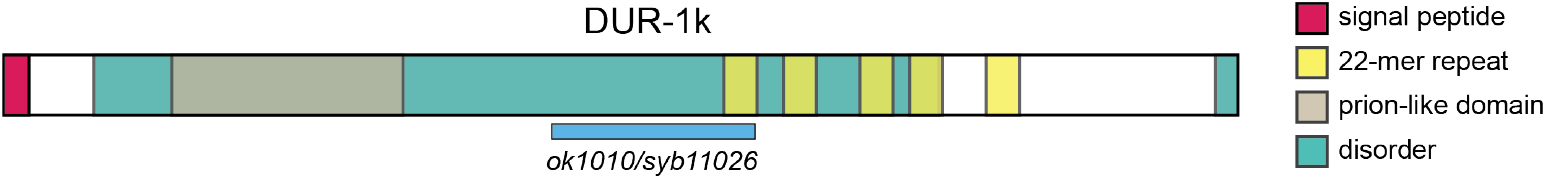
A schematic of the protein domain organization of DUR-1.

